# On the correlation of CTCF binding and c-myc amplification in human prostate cancer cell lines

**DOI:** 10.1101/009761

**Authors:** Tobias T. Fleischmann, Günter Assum

## Abstract

**Abstract:** *Background:* Background: One of the hallmarks of prostate cancer development is the amplification of *C-MYC* copy numbers. It is not yet clear what the driving force behind the activation of *C-MYC* expression in prostate cancer is. CTCF, which is known for its role as chromatin organiser, is thought to be involved in the activation of *C-MYC* expression. The affinity of CTCF to its binding site on the DNA is negatively affected by methylation of CpG islands, including its binding site within the promoter sequences of *C-MYC*. DNA hypermethylation is often found in cancer cells and could therefore be responsible for reduced binding of CTCF at the *C-MYC* promoter, which can in turn lead to enhanced expression.

*Results:* Here, the copy number of *C-MYC* of five different prostate cancer cell lines has been determined by FISH, and the state of methylation of the binding site of CTCF upstream of *C-MYC* was investigated. The *C-MYC* locus was shown to be amplified to various degrees in all cell lines, reflecting the copy number of actual tumors very well, but methylation was found to remain unchanged compared to healthy cells in all examined cell lines. Hence, no correlation between *C-MYC*-copy number and CTCF binding possibilities could be established.

*Conclusions:* The enhanced expression of *C-MYC* in prostate cancer cell lines is not affected by CTCF, whose binding presumably still persists. *C-MYC* activation in prostate cancer must therefore be caused by a disparate mechanism, which also deviates from other well known mechanisms that are seen in Burkitt-Lymphoma, Leukemia and breast cancer.

## 4. Background

*C-MYC* is a very well studied gene, which has been characterised in a plethora of studies during the last decades. *MYC*-gene families are spread all over the vertebrate taxa (1,2,3,4). The human *C-MYC* gene can be transcribed from 4 different alternative promoters, P0, P1, P2 and P3. The prevalent gene product of the *C-MYC* gene is MYC2 (MYC), which is a 439 aa nuclear localised phosphoprotein (5). In its active state, MYC2 is found to be incorporated in a heterodimer together with MAX which shares the same bHLHLZ-Domain as MYC, what allows binding to E-Box-sequences (6,7,8). Another important interaction partner of MYC is BRCA1, which can be downregulated by MYC (9). Furthermore, MYC is involved in the synthesis of certain miRNAs, that are involved either in cancerogenesis (miR-17) or are part of the p53 network (miR-37a) (10,11).

On the genetic level cancer is characterised by widespread DNA damage and deregulation of signal transduction pathways. The key genes involved in carcinogenesis are oncogenes and tumor-suppressorgenes, to which *C-MYC* belongs as a proto oncogen. *C-MYC* appears to play a special role in prostate carcinoma, which is the second most important cause for deadly ending cancer diseases in men in the western hemisphere (12). In prostate cancer amplification and overexpression of *C-MYC* is detectable during almost all stages of tumor development. For example in intraepithelial neoplasies, the predecessors of the actual carcinoma, more than 50% of all tissue samples show an additional chromosome 8, which contains *C-MYC*. In primary prostate carcinoma, anomalies in the *C-MYC* copy number are also found in ˜50% of the tumors, with additional chromosomes 8 as well as translocations of parts of chromosome 8, that include *C-MYC*. In 70% of metastatic carcinoma tissue additional copies are detected with even higher numbers, causing the presence of more than 3 times more *C-MYC* copies than in healthy tissue (13,14). It has been established in tumor samples and also in cell lines, that the copy number is directly proportional to C-MYC protein levels (14,15).

*C-MYC* displays a wide range of regulatory mechanisms by which its expression can be controlled, including a G-Quadruplex structure that is located between the two promoters P0 and P1 and is able to block mRNA-synthesis (16,17,18,19). Another important mode of regulation is of epigenetic nature. This includes phenomena like histone modifications, genomic imprinting, div. chromatin remodeling processes and DNA methylation (20). A common mark, found in tumors, is hypermethylation of stretches of DNA, especially of the promoters of tumorsuppressorgenes, which renders them inactive (21,22). It is evident, that misguided DNA methylation correlates positively with cancerogenesis (23). Misguided methylation can also affect CTCF (ccctc-binding factor), which is an important factor, responsible for the topology and organisation of the whole chromatin structure. CTCF is a 727 aa protein with 11 zink finger domains. It has ˜14000 binding sites in the human genome. One of them is located upstream of *C-MYC*, which also has led to its discovery (24,25,26). Via binding to its target sequences, CTCF is able to activate or repress gene expression by changing chromatin topology. This can bridge long distances and bring enhancers close to genes which normally would not be influenced by them. As the binding of CTCF is methylation dependent, it also takes part in methylation-dependent isolation of eu-from heterochromatin (27,28).

In healthy G0 cells CTCF is constitutively bound upstream of *C-MYC* (29). A possible mode of interaction of CTCF with *C-MYC* would be as a genetic isolator, analogous to its role at the Igf2/H19 locus, where CTCF binds between the promoter and an upstream enhancer element, only in case the CTCF binding site is not methylated and subsequently stops the enhancer from further influencing gene expression (30). This would basically point to a role of CTCF as a repressor of *C-MYC* expression. Another possibility would be that CTCF maintains a barrier and stops euchromatin from spreading into heterochromatin.

It is not known which role CTCF plays in the activation of *C-MYC* that occurs during malignant transformation of cells. It might be possible that upon misregulated changes of methylation in CTCF binding sites of the *C-MYC* promoter CTCF loses its affinity for this particular binding site which in turn leads to enhanced expression of *C-MYC*. As prostate cancer cells usually own more than the usual number of *C-MYC* genes it can also be the case, that different copies of *C-MYC* display different states of methylation at their corresponding CTCF binding site.

The goal of this work was to determine whether there is any correlation between *C-MYC* copy numbers in prostate carcinoma cells lines and an altered methylation status of the CTCF binding site in proximity of *C-MYC*.

## 5. Results and Discussion

### Results

#### Analysis of *C-MYC* copy number in 5 prostate cancer cell lines

The properties of the proto-oncogen *C-MYC*, which promote proliferation, have long been known to the research community. Deregulated expression of *C-MYC* in concert with other factors can lead to cancer. In most of the prostate carcinomas *C-MYC* is detectable in abnormally high copy numbers. Often a simple additional copy of the chromosome 8, which contains *C-MYC*, is found. But in other cases even more copies can arise from translocation events resulting in additional fragments of chromosome 8 on other chromosomes and usually bearing an active copy of *C-MYC*. For the aim of the project it was necessary to determine the genomic background of the cell lines that were used. Therefore all cell lines used in this study have been subjected to a numerical analysis of overall chromosome numbers per cell, numbers of chromosome 8, translocated fragments of chromosome 8 and C-MYC copy number via FISH. On every normal pair of sisterchromatids two copies of C-MYC are present and were counted as one signal, respectively copy, in the genomic context.

#### PC3

For the cell line PC3, derived from bone metastasis of prostate cancer, 29 metaphases were analyzed. On the average the cells showed an overall chromosome number of 56.1 and an average *C-MYC* copy number of 5.6. In 90% of the cases this is composed of 2 normal chromosomes 8 plus 3 to 4 translocated copies of *C-MYC*, exemplified in Figure 1. In this case of 56 chromosomes, one would expect 2.44 copies of a gene, that would normally be present in two copies in an equally amplified genome, if there is no bias of any kind, preferring certain genes. The grade of amplification of *C-MYC* was normalised to the amplified genome and is shown in the column “relative amplification of *C-MYC*” (Table 1). In the case of PC3 the relative amplification of *C-MYC* amounts to an average factor of 2.3.

**Figure 1:**
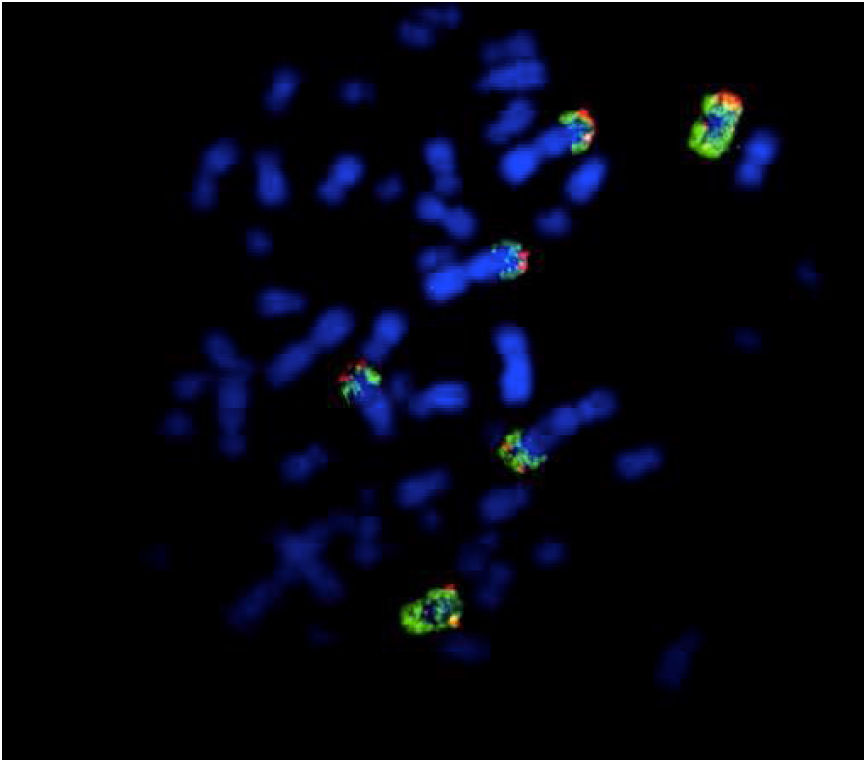
Example for cell line PC3. Blue: 56 DAPI stained chromosomes. Green: 2 Chromosomes 8 and 4 translocations of chromosome 8. Red: Individual red spots indicate one copy of *C-MYC* per sisterchromatid. Two spots on one chromosome have always been counted as one signal during all further analysis steps for all cell lines.

**Table 1:** PC3 shows a merely homogenous distribution of chromosome numbers and additional *C-MYC* copies. The average number of chromosomes per cell is 56 and includes 5 to 6 copies of *C-MYC*, which corresponds to a relative amplification of the factor 2.3.

|  | Chromosome number | Numbers of *C-MYC* | Numbers of chromosome 8 | Numbers of translocations including *C-MYC* | Relative amplification of *C-MYC* |
| --- | --- | --- | --- | --- | --- |
| Group 1 (35 %) | 56 | 5 | 2 | 3 | 2.05 |
| Group 2 (55 %) | 56 | 6 | 2 | 4 | 2.46 |
| Group 3 (10 %) | 57 | 5.5 | 1 | 3.5 | 2.22 |
| Average | 56.10 | 5.60 | 1.90 | 3.60 | 2.30 |

#### 22RV1

With on average 49 chromosomes this cell line of primary prostate carcinoma tissue did not deviate very much from 46 of normal cells. But the average of 3 copies of *C-MYC* displays a overrepresentation of *C-MYC* which is mirrored in its relative amplification of 1.44 (Figure 2 and Table 2). Altogether 30 metaphases were taken into account.

**Figure 2:**
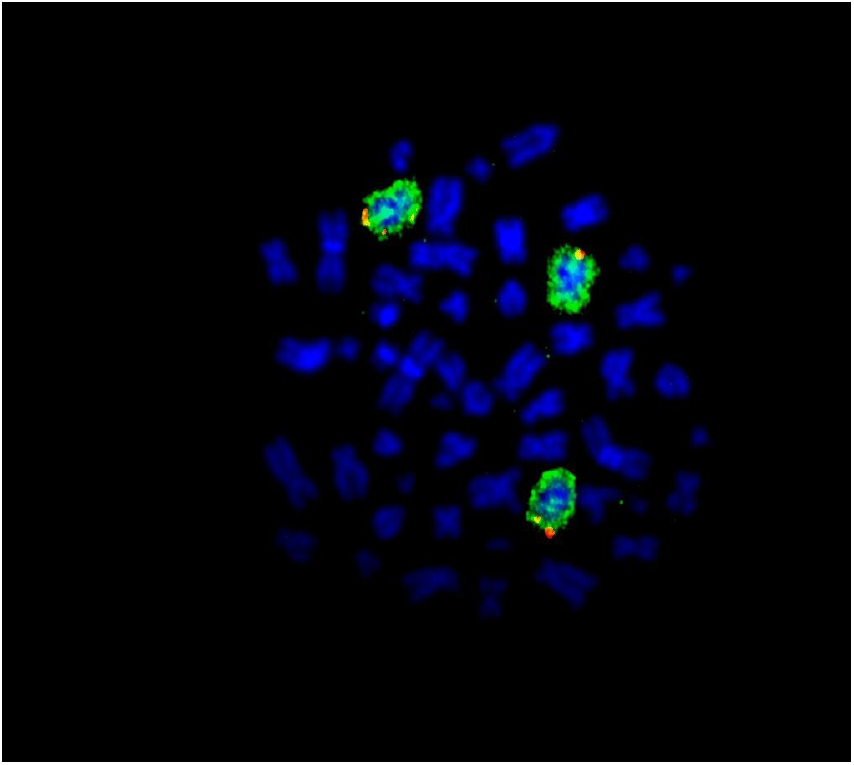
Example for 22RV1. Color code as in Fig.1. The additional copy of *C-MYC* in this case stems from a additional copy of chromosome 8

**Table 2:** 22RV1 shows only a very slight overall amplification of the genome but a significant increase of *C-MYC* copies by the factor of 1.44.

|  | Chromosome number | Numbers of *C-MYC* | Numbers of chromosome 8 | Numbers of translocations including *C-MYC* | Relative amplification of *C-MYC* |
| --- | --- | --- | --- | --- | --- |
| Group 1 (37 %) | 49 | 3 | 3 | - | 1.41 |
| Group 2 (50 %) | 48.5 | 3 | 2 | 1 | 1.42 |
| Group 3 (13 %) | 49.5 | 3.5 | 2 | 1.5 | 1.63 |
| Average | 48.82 | 3.06 | 2.37 | 0.7 | 1.44 |

##### DU145

This cell line, originating from a brain metastasis of prostate cancer, showed the lowest variation of all analyzed lines. On average 60 chromosomes with 3 copies of chromosome 8 are present (Table 3 and Figure 3). 32 cells were subjected to analysis.

**Figure 3:**
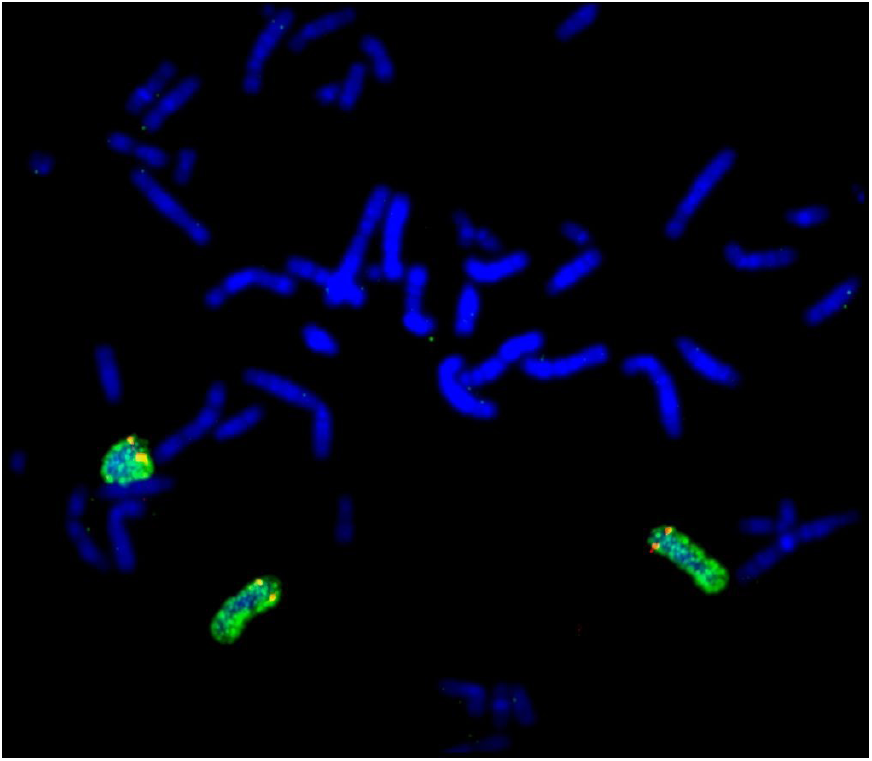
Example for DU145. Color code as in fig.1. Three chromosomes 8 are visible, each containing two single C-MYC loci.

**Table 3:**
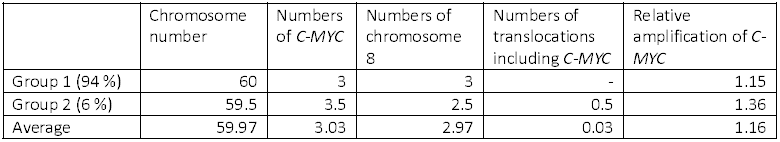
Most of the DU145 cells showed an additional chromosome 8 and very rarely two additional chromosomes 8. The overall amplification is not very high, reaching a factor of 1.16.

##### LNCap

This line, which originates from lymphnode-metastasis of prostate cancer, seems to shatter into 4 subclones mainly determined by increasing overall chromosome numbers (from ˜40 to ˜84) that show a consistent rise in *C-MYC* copy number as well. The average increase of *C-MYC* numbers is 1.24 and is solely caused by additional copies of chromosome 8 (Table 4 and Figure 4). 20 metaphases of this cell line were available and analysed.

**Figure 4:**
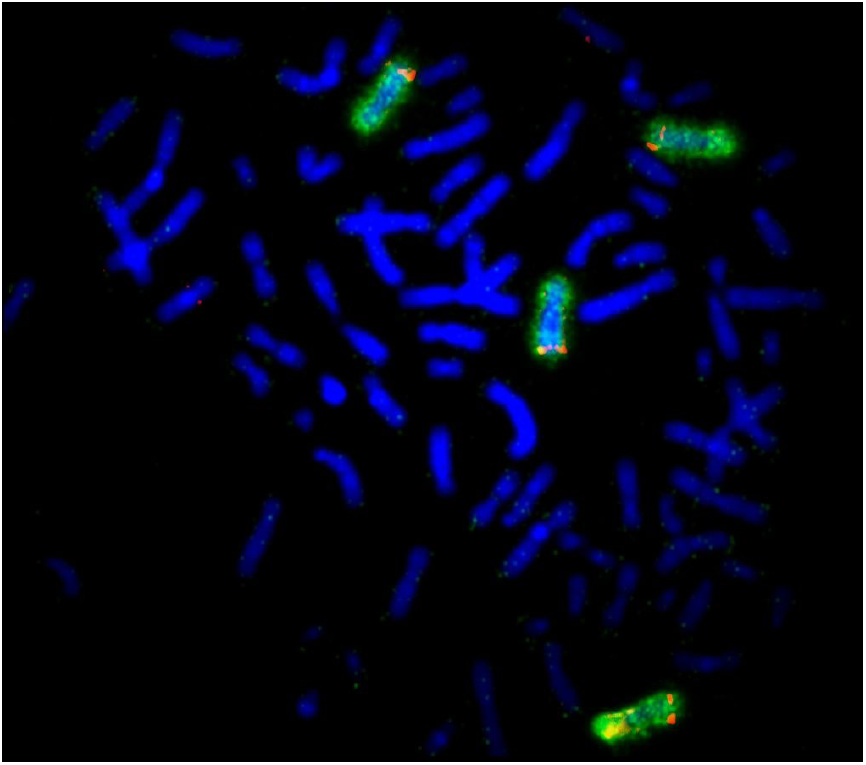
Example for LNCap. Color code as in Fig. 1. Example of LNCap cell with a duplicated chromosome set containing 4 chromosomes 8.

**Table 4:** LNCap displays high diversity regarding its whole genome amplification status. Roughly half of the samples show low chromosome numbers of not higher than 45, whereas the rest almost duplicates the standard chromosome set up to 84. With rising overall amplification there is also a rising number of *C-MYC* copies.

|  | Chromosome number | Numbers of *C-MYC* | Numbers of chromosome 8 | Numbers of translocations including *C-MYC* | Relative amplification of *C-MYC* |
| --- | --- | --- | --- | --- | --- |
| Group 1 (25 %) | 41 | 2 | 2 | - | 1.12 |
| Group 2 (25 %) | 45 | 3 | 3 | - | 1.53 |
| Group 3 (15 %) | 63 | 4 | 4 | - | 1.46 |
| Group 4 (35 %) | 84 | 4 | 4 | - | 1.10 |
| Average | 60.35 | 3.25 | 3.25 | - | 1.24 |

##### PNT1B

Similar to LNCap, this immortalised prostate epithelial cell line is made up of two subclones, that either contain a non-amplified genome or a duplicated chromosome set. The group containing the higher chromosome numbers also tends to have more *C-MYC* copies, mostly as additional copies of the chromosome 8 or in some cases in the form of translocations of chromosome 8 (Table 5 and Figure 5). 31 cells were analysed.

**Figure 5:**
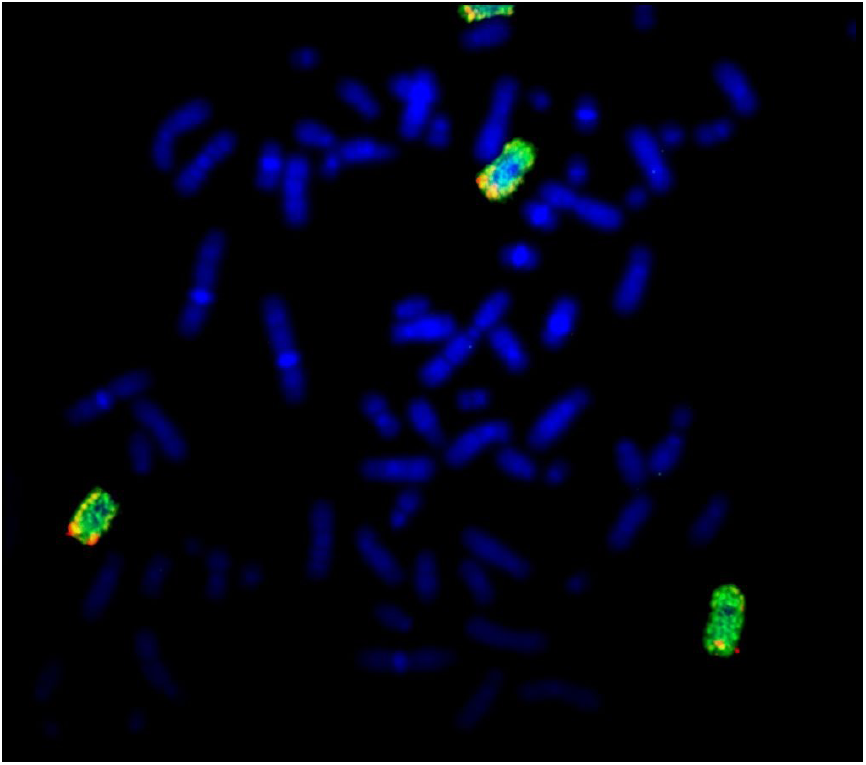
Example for PNT1B. Color code as in Fig. 1. *C-MYC* is slightly overrepresented in the form of additional copies of chromosome 8.

**Table 5:** PNT1B also shows an increased C-MYC number by the factor 1.23.

|  | Chromosome number | Numbers of *C-MYC* | Numbers of chromosome 8 | Numbers of translocations including *C-MYC* | Relative amplification of *C-MYC* |
| --- | --- | --- | --- | --- | --- |
| Group 1 (26 %) | 43 | 2 | 2 | - | 1.07 |
| Group 2 (61 %) | 78 | 4 | 4 | - | 1.18 |
| Group 3 (13 %) | 77 | 5.5 | 4 | 1.5 | 1.64 |
| Average | 68.77 | 3.68 | 3.48 | 0.20 | 1.23 |

#### Summary FISH results

The cell line PC3 owns the highest relative copy number, apparent by the relative amplification of *C-MYC* by 2.3, more than the double amount of *C-MYC* copies that would be expected, if the amplification of the genome would occur randomly. The second strongest amplification, but much lower than in PC3, is represented by line 22RV1. The other three cell lines only display slight increases (Table 6). Students t-test was used to determine P-values.

**Table 6:**
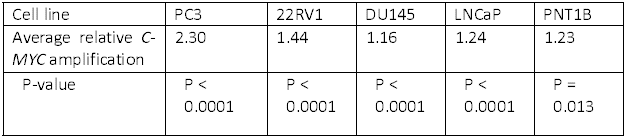
Summary of FISH analysis of all cell lines.

| Cell line | PC3 | 22RV1 | DU145 | LNCaP | PNT1B |
| --- | --- | --- | --- | --- | --- |
| Average relative *C-MYC* amplification | 2.30 | 1.44 | 1.16 | 1.24 | 1.23 |
| P-value | P < 0.0001 | P < 0.0001 | P < 0.0001 | P < 0.0001 | P = 0.013 |

#### Bisulfite sequencing

The amplified *C-MYC* copies, present in most prostate carcinomas, that contribute to a high expression of *C-MYC* are thought to rather be caused by strong upregulation of single copies than low expression of multiple copies. There are mechanisms in the cell that could promote such a strong expression of for example one single copy, but which won’t interfere with the activation of the rest of the copies. One of these putative mechanisms could be carried out by the activity of the genetic isolator CTCF, which is constitutively bound 2 kb upstream of *C-MYC* in healthy tissue. A loss of CTCF’s binding to *C-MYC* could promote uncontrolled expression of *C-MYC*. The binding of CTCF is methylation dependent, therefore methylation of said sequence would in this case disable CTCF binding.

We wanted to test whether the phenomenon of methylated binding sites could be detected in any of the five cell lines. Therefore the methylation status of all CpG islands on a sequence of 300 bp in the area of interest was subjected to bisulfite treatment, subsequent PCR and sequencing. E.g. in a hypothetical cell line with 5 copies of *C-MYC*, two of them could still be under the control of *C-MYC*, whilst the other 3 copies might have methylated CpGs at the CTCF binding site, which could lead to overexpression via access to enhancer sequences. To be able to roughly cover such possible distributions a certain amount of clones is needed to be sequenced. DNA of all lines was bisulfite treated as described and PCR was performed. The products, with the possibility to bear the substituted and unsubstituted nucleotides (corresponding to unmethylated and methylated sequences) were cloned into plasmids and transformed into *E. coli*. 10 to 16 clones of each cell line were subsequently sequenced. No significant change in the methylation status compared to healthy tissue could be discovered. The CTCF binding site, respectively its surrounding region, was not methylated in any of the used cell lines.

#### Discussion

In the first part of this study the copy number of the proto-oncogene *C-MYC* has been determined in prostate and prostate cancer derived cell lines. All cell lines, PC3, 22RV1, DU145, LNCap and PNT1B displayed increases in the number of *C-MYC* genes present in the cells. The increases are not very strong, ranging from a relative enrichment of factor 1.16 to 2.30. This is in contrast to the very high copy numbers found in mouse prostate carcinoma with copy numbers of around 50 (31), but is in line with previous studies performed on human tissues (14,15). In fact this result shows that the cell lines used in this study are reliable model systems for prostate cancers that realistically mirror the situation in genuine cancers regarding *C-MYC* copy numbers. This finding further strengthens the indication that *C-MYC* activation is not driven by the simultaneous expression of multiple copies of *C-MYC* but by a stronger process which is able to interfere with single loci.

One possible candidate for this kind of process is the chromatin-remodeler CTCF, which can block or grant access to enhancers. The closest binding site of CTCF to *C-MYC* contains several CpG dinucleotides, which can be targets of methylation. As the binding of CTCF depends on the methylation status of these islands, said status was determined in this study. The CTCF binding site of *C-MYC* was still unmethylated in all cell lines as it would also be expected for quiescent cells in healthy tissue. Therefore the presence of CTCF on its binding site in the proximity of *C-MYC* is very unlikely to have any effect on the activation of *C-MYC* during malignant transformation of prostate cells. Changes in methylation patterns, especially hypermethylation are often found in various types of cancer, but this locus in particular remains spared of these changes.

## 6. Conclusions

We conclude that in human prostate cancer cells there is no correlation between relative/absolute *C-MYC* amplification and the methylation status of the CTCF binding site of *C-MYC*. Other mechanisms must be responsible for the rise in *C-MYC* expression. Therefore *C-MYC* activation in prostate carcinoma appears to differ greatly in its way of activation from many other types of cancer, e.g. breast cancer and certain cell lines, e.g. HL60 und TD2, where *C-MYC* is found to be amplified to high numbers and is often present in the form of double minute chromosomes or homogeneously staining regions (32,33,34).

## 7. Methods

### Cell lines

Five different cell lines have been used for the experiments: PC 3: bone metastasis, 22rv1: primary prostate carcinoma, LNCaP: metastatic cells of lymph tissue of adenocarcinoma, DU145: brain metastasis of adenocarcinoma, PNT1B: prostate epithelial cells immortalised with SV40 large T Antigen.

### Cell culture

The five adherent cell lines have been cultivated separately in standard culture flasks at 37 °C and 5 % CO_2_ in 1x DMEM medium. The cells have been supplied with fresh DMEM medium three times per week after washing with Hanks medium. After a certain cell density had been achieved the medium was removed and the cells were treated with 3 mL of Trypsin-EDTA solution for 5 minutes at 37 °C to detach them from the flaskwalls. Trypsin was deactivated by dilution to 8.5 mL with 1x DMEM medium. The solution was centrifuged and the pellet was resuspended in DMEM medium. The new cultures were inocculated with the required amount of cells.

### DNA-isolation from cells

Adherent cells were detached from the flask walls and pelleted as described.

The cells were washed in 1x PBS buffer, pelleted and resuspended in 0.5 mL of 1x SE buffer (75 mM NaCl, 25 mM EDTA, pH 8). 5μl of proteinase K and 25 μl of 20 % SDS were added to lyse the cells at 37 ^°^C over night. After digestion 170μl of saturated NaCl solution were added and mixed properly. After a centrifugation step of 15 minutes at 4000 g the supernatant was transfered to new reaction tubes and mixed with 1 mL of 100 % EtOH. The DNA pellet is washed in 70 % ethanol, airdryed and resuspended in 0.5 mL H_2_O.

### Analysis of CTCF binding site methylation status

#### Bisulfite treatment of DNA

DNA from the cell lines was treated according to the instructions of the EZ DNA Methylation Kit/Zymo Research. Freiburg, Germany. After bisulfite treatment the CTCF binding site at the C-MYC promotor was amplified by PCR using the following primers. Forward: 5′-TGAAAGAATAACAAGGAGGTGGCTGGAAA-3′, Reverse: 5′ CTCCTACCTCCAAACCTTTACCACAAACAC-3′, The 295 bp product contains the CTCF binding site. A control PCR for successful bisulfite treatment was performed for all reactions. The size of the PCR product was checked by electrophoresis in a 1.5 % agarose gel. Products of the correct size were cut out of the gel and were extracted using the GFX PCR Purification Kit/GE Healthcare, Buckinghamshire, UK. A small amount of the purified Fragment was again checked on an agarose gel. Subsequently the PCR products were cloned into TOPO vectors using the TOPO TA cloning kit after adding an Adenine overhang at the 3′ ends. (10 μL of the purified and resuspended PCRs product, 2 μL of 20 mM buffer, 0.5 μL Taq-Polymerase, 0.3 μL dATP 20 mM, 10 minutes at 72 °C). The vectors were cloned into competent *E. coli* cells. Clones positive for the used selection markers (Ampicillin, X-gal) were selected for analysis via PCR to distinguish between true and false positive clones. For each plate 24 clones were selected and checked using the primers as before. Plasmid DNA was isolated using the plasmid prep mini spin kit/Qiagen, Hilden, Germany.

### FISH

#### Slide preparation

Culture flasks were treated with colcemide for 15 minutes at a concentration of 100 μg/mL. The cells were spun down for 8 minutes at 1000 rpms and the pellet was resuspended dropwise in 10 mL of 0,4 % (w/v) KCl-solution. After 30 minutes at 37 °C the nuclei were spun down and the supernatant was discarded. Ice-cold fixativum (10 mL, -20 °C/ Methanol-acetic acid: 3:1) was added dropwise and the nuclei were spun down again. This was repeated three more times. For the preparation of the slides three drops were dropped on each slide from a distance of 25 cm. The slides were dried and stored at 4 °C.

#### Probe preparation

Nick-translation was used for the generation of Digoxygenin-11-dUTP labeled C-MYC specific probes following standard protocols. BAC RZPD737H091027D6 was used as probe. The expected fragment length was between 100 and 500 bp.

#### Hybridisation

To free the slides of remaining RNA, they were treated with 150 μL of RNase and were kept for 1 hour at 37 °C. The slides were washed twice in 2x SSC buffer (300 mM NaCl, 30 mM Na-Citrate) for 5 minutes each. To release remaining proteins from the DNA the slides were treated with Pepsine solution (30 μg/mL) for 8 minutes at 37 °C. The Slides were washed 2x for 5 minutes in 1x PBS-Buffer at RT and 1x for 5 minutes in 50 mM MgCl_2_ in 1x PBS. The slides were fixed with 1 % formaldehyde for 15 minutes at RT. Afterwards the samples were dehydrated by treatment with a series of solutions of increasing alcohol content for 3 minutes each, starting with 70 % EtOH, to 80 %, 90 % and 99 %. The slides then were dried for 30 minutes at RT. To denature the DNA strands the slides were held into 70 % formamide in 2x SSC for 30 seconds at 70 °C. The slides were rinsed with 2x SSC at 4 °C and are again dehydrated with the alcohol series as before. The C-MYC probe, as well as the Whole Chromosome Paint 8 (WCP8) were heated to 76 °C for 10 minutes in 50 % formamide solution. 14 μL of each probe were put on each slide, sealed with a thin cover glass and incubated at 37 °C for 2 days. Afterwards the slides were washed in several solutions: Rinse 2x in 2x SSC, then 3x for 10 minutes each at 45 °C in Formamide/4x SSC (1:1). They were rinsed once with 2x SSC, and washed 1x in 2x SSC for 5 minutes and 2x for 5 minutes in 0.2x SSC at 60 °C and stored until the next steps in 4x SSC.

#### Antibody treatment

These steps should be performed in a low light environment. WCP8 is labeled with Biotin-14-dATP and the C-MYC probes with Digoxygenin-11-dUTP. FITC-Avidin was added for the binding to biotin, then Anti-Avidin and again FITC-Avidin. Anti-digoxigenin was bound to Digoxigenin, then TEXAS-Red I and TEXAS-Red II. To avoid unspecific binding the slides were treated with 5 % BSA-solution in 4x SSC with 1 % Tween20 for 30 minutes. At first FITC-Avidin was used at 1:200 together with Anti-Digoxigenin at 1:250 in 1 % BSA-solution. 150 μL per slide were incubated for 30 minutes at 37 °C. The slides were washed with 4x SSC and 1 % Tween20 for 5 minutes each 3x. Antiavidin at 1:100 and TEXAS-Red I at 1:25 was applied as before, and washed. Finally FITC-Avidin at 1:200 and TEXAS-Red II at 1:25 were applied. DAPI staining was done for all slides in the last step. After the slides were dried Anti-fade solution was added.

## 10. Author contributions

T.T.F. has performed experiments, interpreted the data and written the paper. G.A. has designed the research and interpreted data.

